# Evidence for *G6PD* variant classification from multiplexed functional assays

**DOI:** 10.1101/2025.08.11.669723

**Authors:** Renee C. Geck, Melinda K. Wheelock, Rachel L. Powell, Ziyu R. Wang, Daniel L. Holmes, Shawn Fayer, Gabriel E. Boyle, Allyssa J. Vandi, Abby V. McGee, Clara J. Amorosi, Nick Moore, Alan F. Rubin, Douglas M. Fowler, Maitreya J. Dunham

**Affiliations:** University of Washington, Department of Genome Sciences, Seattle, WA, 98195, USA; Bioinformatics Division, The Walter and Eliza Hall Institute of Medical Research, Parkville, VIC, 3050, Australia; Department of Medical Biology, University of Melbourne, Parkville, VIC, 3052, Australia; Department of Bioengineering, University of Washington, Seattle, WA, 98195, USA; Brotman Baty Institute for Precision Medicine, Seattle, WA, 98195, USA

## Abstract

G6PD deficiency is one of the most common enzyme deficiencies worldwide, and increases the likelihood of adverse reactions to certain drugs and foods. Identifying people at risk is challenging, since most are asymptomatic until they encounter a trigger. This is further complicated since over 60% of 1,559 known genetic variants in *G6PD* are variants of uncertain significance and thus cannot guide drug prescribing and dosing. To resolve which variants are clinically meaningful and avoid harm from adverse drug reactions, we conducted two high-throughput functional assays: one for G6PD activity, and one for abundance. We measured the function of 9,527 missense, nonsense, and synonymous *G6PD* variants. The patterns of variant effect on activity and abundance confirmed the importance of structural NADP^+^ for G6PD activity and abundance, and G6PD dimerization for G6PD activity. Based on the ability of our functional assay scores to accurately classify *G6PD* variants of known clinical effect, we generated evidence that 4,870 missense variants contribute to G6PD deficiency and 2,245 are unlikely to contribute to G6PD deficiency. Our data can be used to deepen our understanding of G6PD as a protein, and to close the gap in classification for variants of uncertain significance to improve implementation of genetic medicine for G6PD deficiency.

## Introduction

G6PD deficiency is one of the most common enzymopathies in the world, but most G6PD-deficient individuals have no other symptoms beyond low enzyme activity^1^. However, G6PD deficiency increases the risk of adverse reactions to many pharmaceutical drugs, so identifying individuals with G6PD deficiency is crucial for avoiding these reactions^2^.

G6PD is the main producer of NADPH in red blood cells, which is then used to create antioxidants^3^. In the presence of a trigger, such as fava beans, infection, or certain drugs, insufficient NADPH and antioxidants can lead to lysis of red blood cells and acute hemolytic anemia (AHA)^1^. Because G6PD deficiency increases risk for AHA, G6PD activity levels may be tested before prescribing certain drugs such as rasburicase and primaquine^2^. However, use of testing varies greatly, and test results can be confounded by other hematological parameters^1,4^. Activity tests cannot be used during a hemolytic crisis even if G6PD deficiency is the suspected cause, since the blood cells with the lowest activity will have lysed and thus not be measured, leading to a potential false normal range result.

Genetic testing for G6PD deficiency is attractive because it can be done at any time and may already be available as provider- and consumer-directed genetic testing usage increases. G6PD deficiency is caused by variants within the *G6PD* gene, which is located on the X chromosome^5^. Individuals hemizygous or homozygous for a low-activity variant will have G6PD deficiency, and heterozygous individuals may present with deficiency since their activity can vary due to X-inactivation^1,2^. As of September 2025, 1,559 variants, including 584 single missense variants, have been identified in *G6PD* (https://github.com/reneegeck/G6PDcat)^6^. Of these, 200 missense variants have been classified in ClinVar as “pathogenic” or “likely pathogenic” for their contribution to G6PD deficiency, but over 350 others remain variants of uncertain significance (VUS)^7^. VUS are a major challenge for implementation of genetic medicine, since they cannot be used to inform patient diagnosis or treatment^8^. In order to classify VUS, additional evidence is required, such as information on drug-induced anemia, allele segregation, and functional assays^9^. The World Health Organization (WHO) has also developed a variant classification system specific to *G6PD*, based on median activity measured in at least 3 unrelated individuals. Variants with at least 60% of normal activity are class C, variants with under 45% activity in people with AHA are class B, and variants with under 20% activity in people with chronic non-spherocytic anemia (CNSHA) are class A^10^. All of the class A and B variants change the coding sequence of *G6PD*, supporting that altering the protein sequence of G6PD is the primary molecular mechanism underlying deficiency.

We and others previously surveyed the literature to collect information on *G6PD* variants, leading to a decrease in the number of VUS^2,5,6,11^. However, new variants are frequently reported, often without sufficient data for classification. Functional assays can help close this gap, by directly measuring properties such as enzyme activity and protein stability, or indirectly measuring effects of enzyme activity on growth, which has been done for several *G6PD* variants in microbial systems^5,12,13^. However, waiting to test the function of a variant until it is identified in a person is not rapid enough for point-of-care decisions^5^. Multiplexed assays of variant effect (MAVEs) enable functional measurements of thousands of variants in parallel, so that functional data is preemptively available when variants are identified^14^. MAVEs have been successfully used to classify variants in many pharmacogenes for effect on drug metabolism, and enzyme function for contribution to enzymopathies^15^. In order to provide this level of functional evidence for *G6PD* variant classification, we conducted MAVEs on the coding sequence of *G6PD*, providing functional scores for 8,592 missense variants. We calibrated these scores using variants of known effect so that these scores can be incorporated into variant classification frameworks, and provide evidence that 97 current VUS may contribute to G6PD deficiency.

## Results

We selected two MAVEs for assaying G6PD function: a growth-based activity assay, and a fluorescence-based protein abundance assay (Fig. 1A-B). We directly assayed G6PD function in a *Saccharomyces cerevisiae* yeast system in which NADPH produced by G6PD is necessary for growth. *S. cerevisiae* requires NADPH for synthesis of methionine and antioxidants. When the *S. cerevisiae* homolog of *G6PD*, *ZWF1*, is deleted, the resultant *zwf1Δ* yeast grow poorly in the absence of methionine and presence of hydrogen peroxide, but their growth is rescued by expression of human *G6PD*; *G6PD* variants with reduced function only partially rescue growth^5,12^. We transformed *zwf1Δ* yeast with a barcoded plasmid library containing human *G6PD* variants (median 6 barcodes per variant) (Table S1), and used continuous culture turbidostats^16^ to grow them in media lacking methionine with 0.25mM hydrogen peroxide to select for yeast expressing functional *G6PD* variants (Fig. 1A). We also conducted a control in media with methionine and without hydrogen peroxide. Variant barcodes were sequenced from samples taken at ten time points over 72 hours (Table S2) and used to calculate an activity score with a linear regression model, and scaled so the average of all synonymous variants scores was 1 and nonsense variants 0 (see Methods). The common *G6PD* variants A (c.376A>G), which is not associated with deficiency, and Mediterranean (c.563C>T), which is well established to cause deficiency, behaved as expected (Figure S1B)^3,5^.

**Figure 1:**
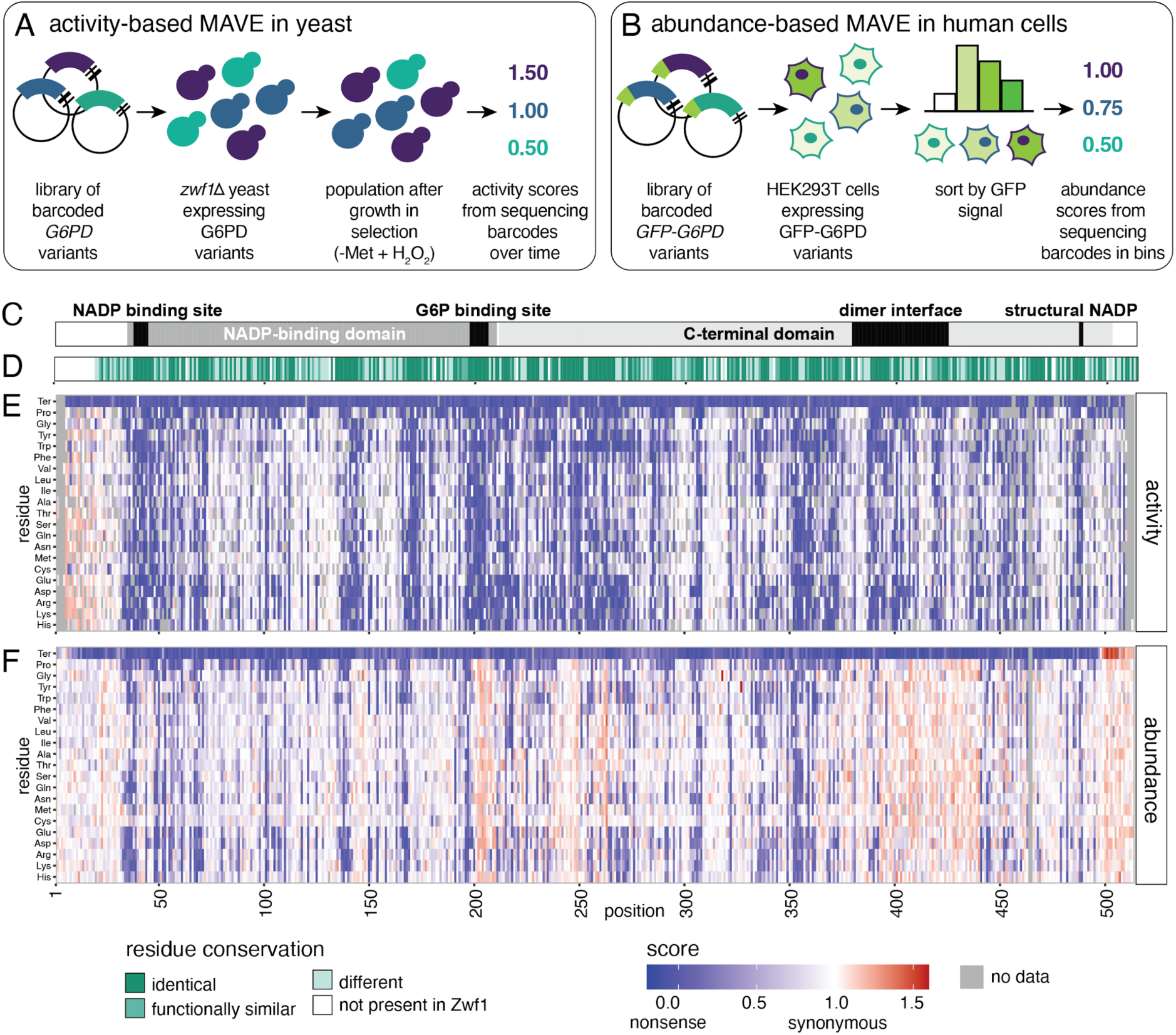
Mutational scans of G6PD for activity and abundance. (A) Schematic^15^ of yeast-based activity assay and (B) VAMP-seq. (C) G6PD domains^18,22^. (D) Conservation of residues from *S. cerevisiae* Zwf1 to *H. sapiens* G6PD. (E) Activity score map of G6PD from yeast-based activity assay in selective condition. Scores are determined from two replicates. (F) Abundance score map from VAMP-seq. Scores are determined from two replicates.

We computed activity scores for 8,228 missense, 481 nonsense, and 435 synonymous variants from growth in the selective condition (Fig. 1C-E, Table S3). This is 84.7% of all 10,794 possible single *G6PD* variants excluding the start and termination codons, comprising 84.3% of 9,766 possible single missense variants. Scores correlated well between replicates in selection (R^2^=0.952, Fig. S1C), and nonsense and synonymous variants were well-separated (Fig. S1D). Variants that increased G6PD activity were rare, with only 25/8228 (0.3%) scoring over 20% higher than the average synonymous variant. The distribution of synonymous scores had a long tail (Fig. S1D), but all synonymous variants did increase in frequency over time in selection (Fig. S1E), as expected when variants with increased activity are rare. All nonsense variants decreased in frequency over time, as expected, except for p.Ser40Ter (Fig. S1E); this may be due to high read-through of the UAG-G sequence in *S. cerevisiae*^17^. About half of missense variants (4104/8228, 49.9%) had activity below 45% of synonymous, which is the cutoff for deficiency set by the World Health Organization (WHO)^10^. Variation at highly conserved residues and in critical functional domains such as the NADP^+^ and glucose-6-phosphate (G6P) binding sites and parts of the dimerization domain decreased activity, as expected (Fig. 1C-E)^18,19^.

We also conducted a MAVE for G6PD abundance. Loss of stability—and thus decreased protein abundance—is a major mechanism contributing to G6PD deficiency, especially for very low function variants that can lead to CNSHA. To interrogate the effects of variants on G6PD abundance, we conducted variant abundance by massively parallel sequencing (VAMP-seq) (Fig. 1B)^20^. The assay was performed by recombining a barcoded GFP-tagged *G6PD* variant library into HEK293T cells (see Methods, Table S1)^21^, with each variant mapping to a median of 12 barcodes. The ratio of GFP to a co-transcriptionally expressed mCherry was used to determine the relative abundance of G6PD variants independent of differences in transcript level. Cells were sorted into four bins based on the ratio of GFP to mCherry fluorescence intensity, and the barcodes in each bin sequenced to determine relative abundance of each variant. As with the yeast activity assay, abundance scores were normalized so the average of all synonymous variants was 1 and nonsense variants 0. Replicates correlated well (R^2^=0.717), with most abundance scores clustered near the synonymous or nonsense values (Fig. S2).

We computed abundance scores for 9,677 missense, 508 nonsense, and 489 synonymous variants, covering 89.7% of possible single missense variants in *G6PD* (Table S3). As expected, variants altering residues 488-489, which interact with the structural NADP^+^ molecule, led to decreased abundance (average of all missense scores at each site 0.3393 and 0.4044, respectively) (Fig. 1F)^19,22^. Many variants in the G6P binding site or dimerization domain had normal or increased abundance, though those variants also had low activity (Fig. 1E-F).

To compare variant activity and abundance, we set thresholds for “low” and “normal” based on the distribution of the synonymous variants, with normal above a z-score of -1 and low below a z-score of -3. Most *G6PD* variants with low abundance in the VAMP-seq assay also had decreased activity in the growth-based activity assay (Fig. 2A, lower left quadrant), except for nonsense variants that truncated the protein within the last exon and led to increased abundance (Fig. 1F). This score distribution is similar to patterns observed for other proteins^23^. Many of the low-abundance, low-activity variants mapped to buried residues in alpha helices and beta sheets, or interacted with NADP^+^ molecules at residues 38-44 and 488-489 (Fig. 2B-C)^18,22^. Other variants decreased activity but not abundance, and primarily affected residues near the dimer interface, and in the substrate binding sites (Fig. 2B). Dimerization or tetramerization is necessary for G6PD activity^19,24^, and we observed many substitutions within the dimerization domain decreased activity (Fig. 2B-C, Fig. 1E). Some substitutions in the dimer interface, particularly to residues without hydrophobic side chains, have been shown to reduce protein stability, but our results suggest that most substitutions at the dimer interface do not greatly decrease total abundance of G6PD protein even if they may disrupt stability or formation of the dimeric form^25^.

**Figure 2:**
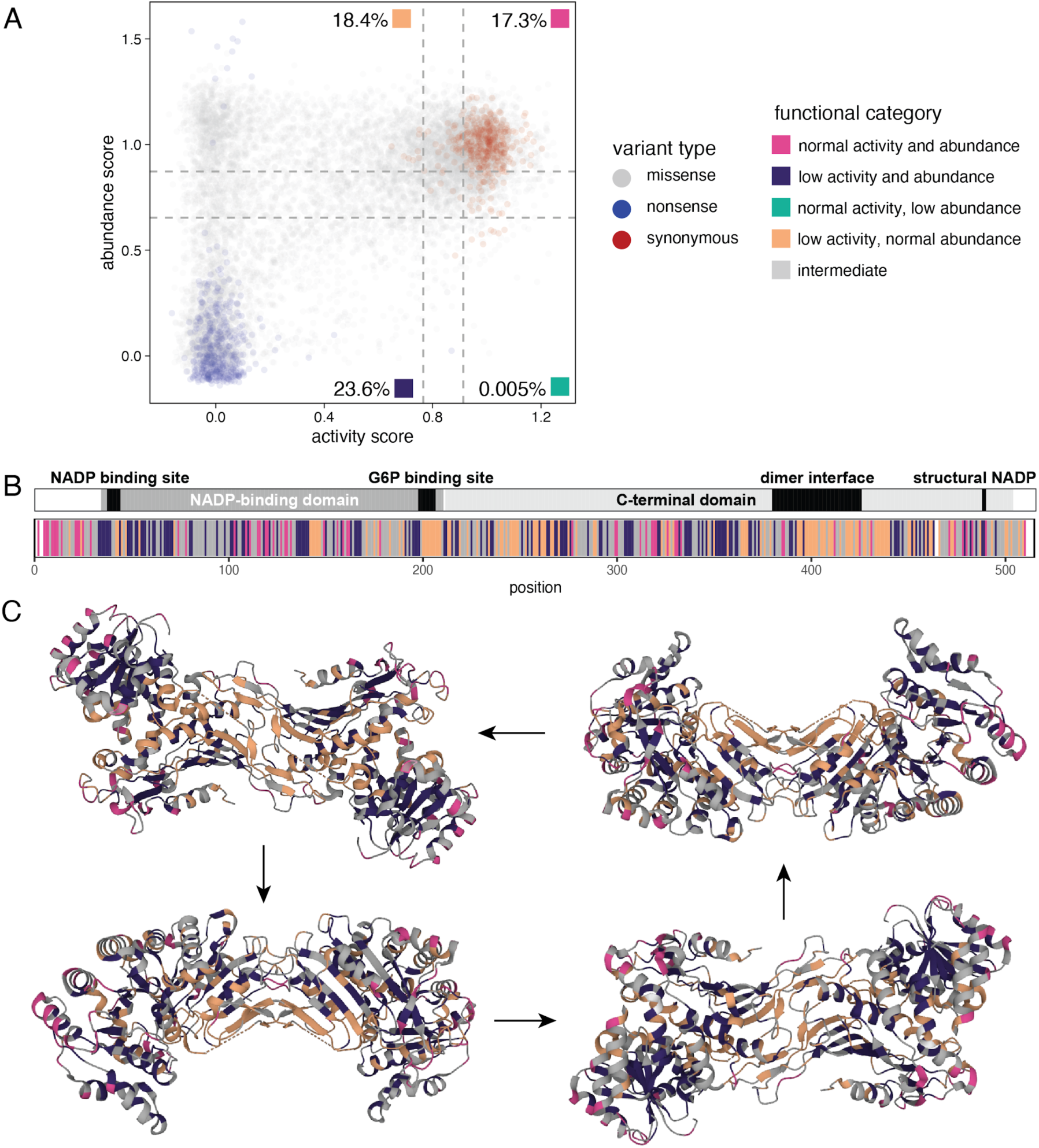
Patterns in variants disrupting activity and abundance. (A) Activity score plotted against abundance score for each variant. Average of all synonymous is set to 1, and average of all nonsense set to 0. (B) Location of low abundance / low activity and low activity / normal abundance variants on G6PD sequence and (C) G6PD homodimer structure (7UAG). Each arrow indicates a 90° rotation towards the viewer around a horizontal axis.

To maximize the utility of our scores, we combined activity and abundance scores into one functional score (Fig. 3). In line with previous strategies for combining MAVE scores^26^, we selected the lower of the activity or abundance score as the functional score because a false normal would have more potential to lead to patient harm by obscuring the presence of G6PD deficiency and risk for adverse drug reactions. The exception was for variants where there was no activity score; variants without activity data only received a functional score if the abundance score was low, indicating a likely loss of function due to low abundance. Variants with normal or intermediate abundance could have normal or reduced catalytic activity, so an abundance score alone was considered insufficient evidence of normal function. We computed these functional scores for 9,527 variants.

**Figure 3:**
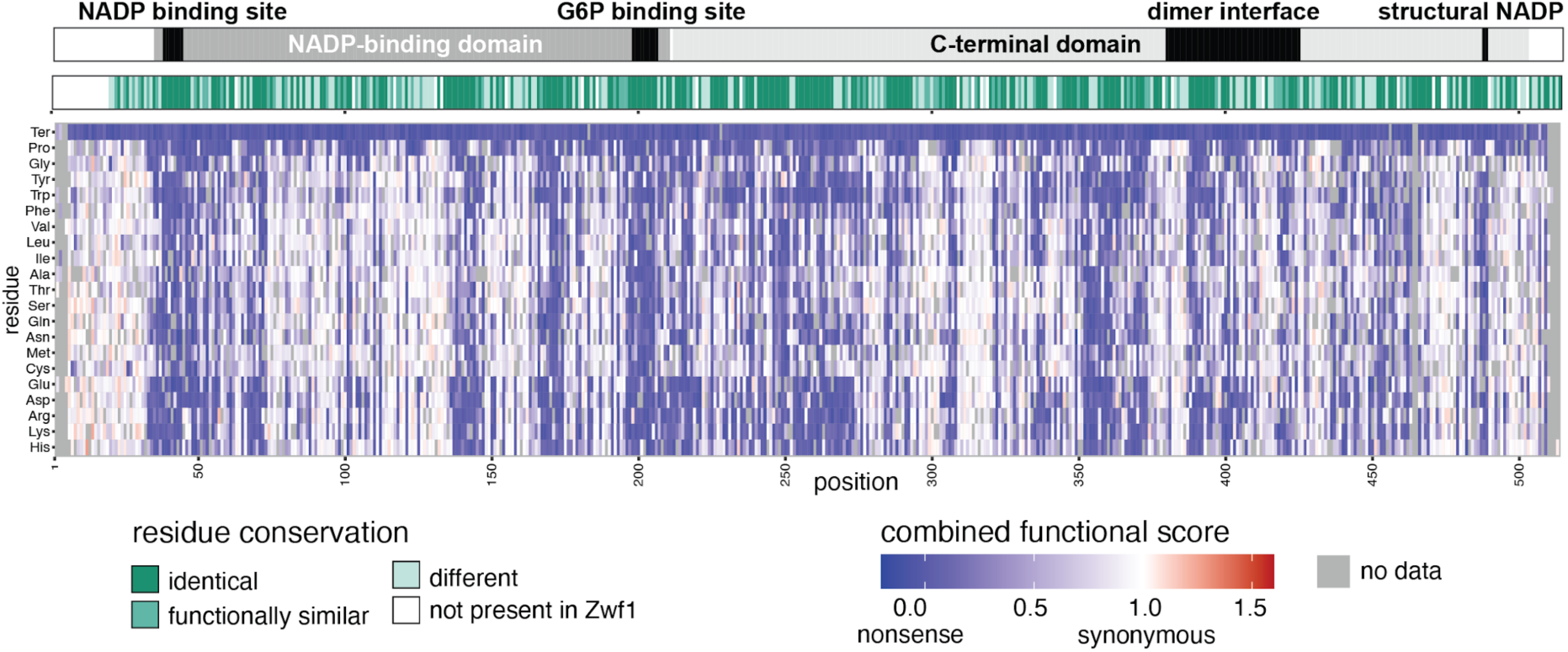
Functional scores of G6PD variants. Heatmap of functional scores. Domains and conservation as in Fig. 1C-D.

When compared to associated phenotypes, variants reported with CNSHA had the lowest functional scores (Fig. 4A). Comparison to WHO classifications (Fig. 4B) was hindered by the small number of variants in Class A: since these variants are rare, few meet the classification requirement of measurements from three unrelated individuals. However, all functional scores below 0.5 are associated with pathogenic variants or variants of uncertain significance on ClinVar. All the variants classified as normal - “C” by WHO or “benign/likely benign” by ClinVar - have normal-like functional scores in our assay (Fig. 4B-C). We observed similar trends with variant effect predictors (VEPs) REVEL and AlphaMissense, where variants predicted to be deleterious had lower functional scores (Fig. S4A-B). Since VEPs and functional data are both considered as evidence for variant classification^9^, adding functional data greatly increases the amount of evidence available for *G6PD* variant classification.

**Figure 4:**
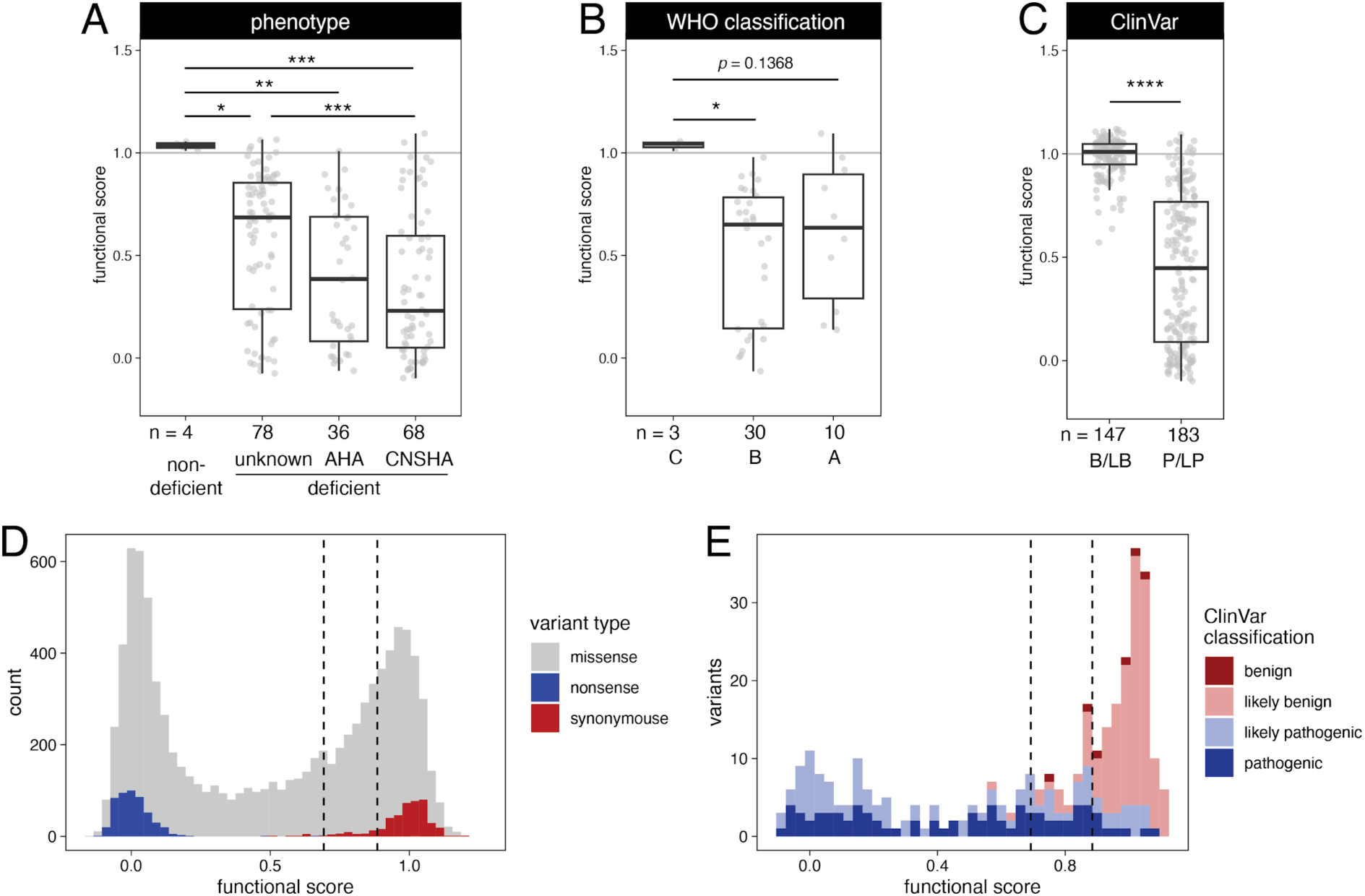
Variant scores as functional evidence for interpretation. (A) Functional scores for variants by phenotype, (B) WHO class, and (C) interpretation on ClinVar: B/LB indicates benign or likely benign, P/LP pathogenic or likely pathogenic. (D) Distribution of variant functional scores. Vertical dashed lines represent z-scores of -1 and -3 based on the synonymous variant distribution. (E) Functional scores of variants classified on ClinVar as known pathogenic / likely pathogenic and benign / likely benign. Significance from by ANOVA with Tukey’s HSD (A-B) or t-test (C); *, p<0.05; **, p<0.01; ***, p<0.001; ****, p<0.0001.

The guidelines for variant interpretation from the American College of Medical Genetics (ACMG) incorporate evidence from functional assays into their scoring system^9^. To determine the weight of evidence given to different scores by a functional assay (increasing in weight from supporting to moderate to strong to very strong), scores must be calibrated by variants known to be pathogenic or benign^27^. Based on the synonymous variant functional score distribution, we used a z-score of -1 as the minimum score for benign evidence, and a z-score of -3 as the maximum score for pathogenic evidence. Applying an OddsPath analysis on variants with interpretations for G6PD deficiency of pathogenic or benign available on ClinVar, we determined that functional scores above 0.8848 can be used in the ACMG framework at up to moderate evidence that the variant is benign (BS3_M, OddsPath = 0.1579), and functional scores below 0.6923 as up to strong evidence that the variant is pathogenic (PS3, OddsPath = 51.01)^27^. In total, 4,870 missense variants were given evidence for pathogenicity; 226 of these variants have been previously reported as found in people, including 105 VUS. 2,245 missense variants were given evidence that they may be benign, including 192 found in people, 167 of which are VUS. This information will be crucial for ongoing *G6PD* variant classification and interpretation.

## Discussion

We present a set of scores for the function of *G6PD* variants, including measurements of both activity and abundance. With increasing numbers of variants identified, functional evidence is critical for addressing the challenge of variants of uncertain significance^8^. Classifying VUS is key to increasing the utility of genetic testing results for G6PD deficiency, which are particularly important in heterozygotes, patients with potentially confounding hematological conditions, and in cases where prescribing decisions can be made rapidly by consulting preexisting test results.

Our functional scores for 9,527 variants, including 8,592 missense variants, correlated well with the known function of many well-characterized *G6PD* variants. Of 161 benign or likely benign variants on ClinVar, we have additional evidence to support that classification for 128/147 that were measured in our assays. For only two likely benign variants do our scores provide conflicting evidence that they may instead be contributing to deficiency (scores of 0.57-0.64); these are both synonymous variants, so the lower scores are likely a result of the long tail of our synonymous distribution (Fig. S1D-E). Of 197 pathogenic or likely pathogenic variants on ClinVar, we have additional evidence to support that classification for 127/183 measured. For 24 we have conflicting evidence that they may not be contributing to deficiency; of these, only eight have multiple submissions to ClinVar so we are more confident that they do contribute to deficiency, and sought to understand why our MAVEs may not be able to measure their effects on G6PD activity or abundance. We noted that residues 374-387 on average have higher functional scores than surrounding residues, despite seven variants associated with chronic anemia falling in this region. These residues are just before and at the beginning of the dimerization domain, so our assays may not be fully capturing the effects of G6PD dimer formation on activity. Overall, the high concordance between clinical classification and our MAVE data support the utility of our scores as functional evidence to aid in classification.

Our data also highlight areas of potential interest for future studies on G6PD structure and function. While we identified very few variants with both increased activity and abundance, many missense variants within N-terminal residues moderately increased activity in our yeast assay. This may be unique to the yeast system, since residues 1-19 are not found in the yeast homolog Zwf1 (Fig. 1D-E); further activity tests in human cells could address this finding. We also observed missense variants in some regions, specifically the G6P binding site, dimer interface, and c-terminus, that increased abundance, and could be informative for structural analysis of the protein (Fig. 1F). G6P binding stabilizes prokaryotic G6PD proteins which lack the structural NADP^+^ that stabilizes eukaryotic G6PD^29^, so our data may support G6P also stabilizing eukaryotic G6PD. Several sites within the dimerization domain were enriched for variants that decreased activity, supporting the importance of recently developed small molecules that stabilize the dimer complex, increase G6PD activity, and may have therapeutic applications for treating G6PD deficiency^19,28,30^.

While most G6PD variants that contribute to deficiency are single missense variants, other types of variants have been identified: in-frame deletions of 1-8 amino acids, and haplotypes comprising 2-3 missense variants, including the common variant G6PD A−, p.[Val68Met;Asn126Asp]^3,6^. Conducting additional functional tests, or MAVEs in the context of different haplotypes, could enable classification of more complex *G6PD* variants. Screening variant libraries in the presence of different G6PD inhibitors and activators could also provide further insight into protein structure and function, and prioritize which variants are most likely to have functionality restored through activator treatment^13,19,28,30^.

More broadly, we highlight important lessons for the growing field of MAVEs. Continuous culture systems are increasing in popularity for selection in growth-based MAVEs^31–33^, and we note that even in the absence of additional selection we could identify nonsense-like variants by their slightly reduced growth in control conditions (Fig. S1D). Conducting a MAVE at different strengths of selection—for instance, increasing the concentration of hydrogen peroxide—could enable better identification of variants with intermediate activity. Repeating the assay with increased selection could address some discrepancies between our functional scores and reported effects of variants, where some variants associated with G6PD deficiency did not receive low functional scores (Fig. 4E).

Guidelines for incorporating functional assay data into variant classification are critical for clinical implementation of these scores^34^. Important work has been done to create general guidelines for their use within the ACMG framework^27^. However, especially with bespoke assays connected to the specific function of a gene product, input from experts on the function and clinical presentation is invaluable. Ongoing work by Variant Curation Expert Panels, such as one specific to *G6PD*, will further guide the specific application of functional scores (https://clinicalgenome.org/affiliation/50147/). With the current score cutoffs we have set, we provide evidence for 272 current VUS: that 105 may contribute to G6PD deficiency, and that 167 are unlikely to contribute. In total, we computed functional scores for 8,592 missense variants, with 4,870 as evidence for pathogenicity and 2,245 for benignity, which will aid in variant classification as more variants are found through case studies and sequencing efforts.

Our data are a resource both for classification of *G6PD* variants and the development of future MAVEs. Measuring the function of pharmacogene variants at scale is key to understanding structure-function relationships and for classification of the rising number of VUS.

## Material and methods

### Strains, cells, and reagents

Yeast lacking *ZWF1* (strain YMD4438, *zwf1::KanMX ura3-HIS3 LEU2 LYS2*) were dissected from the heterozygous diploid deletion collection and phenotyped previously^5,35,36^. HEK293T cells were acquired from ATCC and cultured as described below.

### Library synthesis and barcoding

A pooled dsDNA single site variant library of *G6PD* codons 2-515 was ordered from Twist Biosciences based on the coding sequence of NM_001042351.3 (encoding NP_001035810.1), with flanking homology to pKAM10-LCS (Table S1). One codon was used for each variant, based on codon preference in humans. QC performed by Twist reported successful generation of 10,143 variants (excluding the WT reference sequence), with residues 437 and 464-465 failing synthesis.

For expression in yeast, the oligo library was amplified for 10 cycles to produce homology to yeast expression vector p416-CYC1, which contains the *S. cerevisiae CYC1* promoter and terminator, AmpR for bacterial selection, and *URA3* for yeast selection (Table S1)^37^. Amplicons of the correct length were size selected by adding 0.5 volumes of AMPure XP beads (Beckman Coulter A63881) followed by two 95% ethanol washes, 15 minutes incubation with PEG8000-0.75M NaCl, and two more ethanol washes^38^. p416-CYC1 was digested using BamHI-HF and SalI-HF (NEB R3136 and R3138) and vector and insert assembled in a 1:3 ratio in Gibson Assembly Master Mix (NEB M5510). Any unassembled vector was digested with SalI-HF and the reaction cleaned with DNA Clean & Concentrator-5 (Zymo D4004). The resulting reaction was transformed into electrocompetent NEB10-beta *E. coli* cells (NEB C3020), which were recovered for 1 hour in supplied outgrowth media and transferred to grow in 50mL LB+Amp overnight at 37°C. Variant library plasmids were extracted using ZymoPURE II Plasmid Midiprep Kit (Zymo D4200). For barcoding, the variant library was cut with EagI-HF (NEB R3505). Oligos were ordered from IDT with 18 random nucleotides flanked by homology to the cut site. Gibson assembly with 1:10 vector:insert ratio, re-cutting with EagI-HF, and electroporation was performed as above with the final recovery volume split between four dilutions in 50mL LB+Amp (1:100, 1:200, 1:400, 1:800), which were grown overnight and midiprepped. The library closest to 250,000 unique barcodes based on barcode sequencing was selected for use in the experiments.

For GFP tagging and expression in human cells, pKAM10 with a modified multiple cloning site (pKAM10-LCS), which contains a N-terminal eGFP tag (Table S1), was cut with EcoRI-HF and SpeI-HF (NEB R3101 and R3133) and assembled with the *G6PD* library produced by Twist Biosciences; Gibson assembly followed by re-digestion with EcoRI-HF and electroporation were performed as above. Variant library plasmids were extracted using GenElute HP Plasmid Midiprep Kit (Sigma Aldrich NA0200). For barcoding, the variant library was cut with NotI-HF and XmaI (NEB R3189 and R0180) and assembled with random barcode oligos generated by PCR from the barcode oligos used in the yeast library (Table S1). Electroporation and library selection were performed as above.

### Generation of barcode-variant maps

The barcoded yeast expression plasmid library generated in *E. coli* was linearized with SphI-HF (NEB R3182) and and prepared using SMRTbell Express Template Prep Kit 2.0 (Pacific Biosciences 100-938-900) according to manufacturer’s instructions, followed by nuclease clean-up and an additional cleaning with 0.45 ratio of AMPure PB beads. The barcoded GFP-tagged human expression plasmid library generated in *E. coli* was digested with MluI-HF (NEB R3198) and prepared using the SMRTbell kit with 0.6 ratio of AMPure PB beads.

Samples were submitted to University of Washington PacBio Sequencing Services; the yeast expression library was sequenced on Revio v1 SMRT Cell 25M with 24 hour movie time and the human expression library sequenced using 50% of reads from a Sequel II v3.1 SMRT Cell 8M with 30 hour movie time. Revio reads were filtered using bamtools for read quality over 0.999, and Sequel II filtered using samtools for reads with at least three passes. Barcode-variant maps were generated using Pacybara (February 15, 2024 release) with recommended settings for Revio (MINBCQ=32, MINQUAL=27) or Sequel II data (MINBCQ=62, MINQUAL=93)^39^.

### Activity-based DMS in yeast

Yeast lacking *ZWF1* (*zwf1Δ,* strain YMD4438) were transformed by a high-efficiency lithium acetate / carrier DNA / PEG method^40^. Yeast were grown overnight and backdiluted to OD_600_=0.45 in 50mL YPD, grown for ∼4 hours to reach OD_600_=1.8, washed, and transformed in ten sub-reactions with 1µg plasmid library each. Transformants were recovered for 1 hour in YPD + 0.5M sorbitol at 30°C, pelleted, and resuspended in 50mL sc-ura for selection of *URA3*-marked plasmids overnight at 30°C. Ten stocks of the library were saved for experiments to avoid freeze-thaw cycles or multiple outgrowths.

The *zwf1Δ* yeast library expressing barcoded *G6PD* variants was grown overnight in sc-ura at 30°C. Turbidostats were assembled as previously described^16^, with two containing sc-ura (control) and two containing sc-ura-met + 0.25mM H_2_O_2_ (selective); uracil was omitted in all conditions to maintain the *URA3*-marked plasmids. Yeast were inoculated at OD_600_=0.25 in 200mL per turbidostat, half of the setpoint OD_600_=0.5, and grown at 30°C. An initial sample was taken from the inoculant, and nine additional samples were collected from the effluent at indicated times over 72 hours (Fig. S1A); more samples were taken within the first 24 hours intending to capture variants with very low activity that were rapidly lost from the population. For effluent collection, three 3mL samples were taken from each turbidostat per time point for redundancy, centrifuged at 7000 x g for 3 minutes, and pellets stored at -20°C. OD_600_, total pump run time, and effluent volume were also measured at each collection time point. Number of population doublings at each time point was calculated based on pump run time, since 0.252mL of media are added every 1 second the pump is on. The 72 hours of the experiment was equivalent to 30.3-30.6 population doublings in the two replicate selection turbidostats and 34.8-36.5 population doublings in the controls (Table S4).

DNA was extracted from harvested yeast pellets using Zymoprep Yeast Plasmid Miniprep II (Zymo D2004). Barcode regions were amplified using KAPA2G Robust (Roche KK5702) for 5 cycles with 60°C annealing temperature and 15 seconds extension (Table S1). Samples were cleaned using DNA Clean & Concentrator-5 (Zymo D4004), and the entire reaction used to add indices and Illumina adaptors by an additional 10 cycles of KAPA2G PCR with 65°C annealing temperature and 15 second extension (Table S1). Samples were cleaned again, eluted in 10µL water, and concentration determined on a Quantus fluorometer (Promega E6150) using Quant-iT dsDNA HS Assay Kit (Invitrogen Q32851). Equal amounts of samples were pooled, cleaned using 1.8 ratio of AMPure XP beads, and sequenced on an Illumina NextSeq 550.

### VAMP-seq

The human variant library was transfected into HEK293T cells expressing a Tet-on landing pad (HEK293T TetBxb1BFP Clone4)^20^ concurrently with the Bxb1 recombinase-expressing pCAG-NLS-Bxb1 plasmid (Addgene #51271) using FuGENE 6 (Promega) according to the manufacturer’s protocol. Two days following transfection, variant expression was induced by adding 2μg/mL doxycycline to the media (DMEM + 10% FBS + 1% penicillin-streptomycin). Four days after transfection, cells were treated with 10nM rimiducid (AP1903) to selectively kill unrecombined cells. One day following treatment with rimiducid, media (DMEM + 10% FBS + 1% penicillin-streptomycin) and 2µg/mL doxycycline were replaced. Two to four days after treatment with rimiducid, depending on culture confluence (90-100%), cells were prepared for sorting by lifting from plates with Trypsin-EDTA 0.05% (Gibco), quenching with media, spinning at 300 x g for 5 minutes, aspirating the supernatant, resuspending the cell pellet in sort buffer (1X PBS + 0.5% BSA), and filtering through 35um nylon mesh.

Cells were sorted on a BD Aria III FACS machine using an 85µm or 100µm nozzle. Live cells were first identified using FSC-A vs. SSC-A. Live cells were then gated for single cells using two sequential gates: FSC-A vs. FSC-H and SSC-A vs. SSC-H. mTagBFP2 expression from the unrecombined landing pad was excited using the 405nm laser and captured on the 450/50 nm bandpass filter. mCherry, expressed from successful recombination of the library plasmid, was excited with a 561 nm laser, and emission was detected using a 600 nm long pass and 610/20 band pass filters. EGFP, expressed after successful recombination of the library plasmid, was excited with a 488nm laser and captured using a 505nm long pass and 530/30nm band pass filters. Recombinant mTagBFP2-negative, mCherry-positive cells were isolated, with mCherry fluorescence values at least 10 times higher than the median fluorescence value of negative or control cells, and mTagBFP2 fluorescence at least 10 times lower than the median of the unrecombined mTagBFP2 positive cells. mCherry-positive cells were then gated on EGFP fluorescence, and sorted into four approximately equally sized quartile bins. All flow cytometry data were collected with FACSDiva v.8.0.1 and analyzed using FlowJo v.10.10.0.

After sorting, each bin of library cells was spun down at 300 x g for 5 minutes, then re-plated in 10mL cultures with media and 2μg/mL doxycycline. Cells were allowed to grow until 90-100% confluence. To harvest, cells were lifted from plates with Trypsin-EDTA 0.05%, quenched with media, counted via hemocytometer, then spun at 300 x g for 5 minutes. The supernatant was aspirated, and the pellet was flash frozen in liquid nitrogen before storage at - 20°C.

Genomic DNA was extracted from harvested cell pellets using the DNeasy Blood and Tissue kit (Qiagen) according to the manufacturer’s protocol with the addition of a 30 minute RNase digestion at 56°C during the resuspension step. Each cell pellet was split into multiple DNeasy columns to avoid overloading the column (approximately 5 x 10^5^ cells per column). The human G6PD with barcode region was amplified from genomic DNA using Invitrogen Platinum SuperFi II PCR Master Mix (Invitrogen 12368010) for 5 cycles with 65°C annealing temperature and 120 seconds extension (Table S1). Samples were cleaned using a 0.5 ratio Cytiva Sera-Mag Select beads (Cytiva 29343052) following the manufacturer’s instructions. The barcode region was amplified from the cleaned product using KAPA2G Robust (Roche KK5702) for 5 cycles with 60°C annealing temperature and 15 seconds extension (Table S1). This second PCR product was cleaned using a 2.5 ratio Cytiva Sera-Mag Select beads following the manufacturer’s instructions. The cleaned product was used to add indices and Illumina adaptors by an additional 10 cycles of KAPA2G PCR with 65°C annealing temperature and 15 sec extension (Table S1). Samples were cleaned again, eluted in 10µL, and concentration determined on a Qubit Fluorometer 2.0 using Qubit dsDNA HS Assay Kit (Invitrogen Q33231). Equal molar amounts of samples were pooled, cleaned using 1.8 ratio of AMPure XP beads, and sequenced on an Illumina NextSeq 2000 using a P3 50-cycle kit.

### MAVE data analysis

Sequence files in FASTQ format were produced by bcl2fastq v2.20 and then merged using PEAR^41^ v0.9.11 with minimum overlap, minimum assembly length and maximum assembly length all set to the barcode length of 18. Barcodes were mapped to sequences using the output of Pacybara (described above), with sequences mapped to protein variants using the protein variant caller in CountESS v0.0.79 (https://zenodo.org/records/16357955). Barcodes and sequences of incorrect length, and protein variants represented by less than three distinct barcodes, were removed from the analysis.

For the activity-based MAVE in yeast, barcodes were mapped from 37 assembled FASTQ files, with one file representing the input library and four sets of nine files representing the two control and two selection replicates at times 4, 8, 12, 16, 24, 36, 48, 60 and 72 hours. Counts were summed per protein variant per sample, and expressed as a fraction of the total count for the sample. Variants were scored using a linear regression (ordinary least squares) of their fraction versus the dilution volume at the time of the sample, with raw score determined from the coefficient. Higher activity variants have a positive coefficient as their population grows, whereas lower activity or nonfunctional variants have a negative coefficient as their population dwindles. Raw scores were then normalized by scaling so the average of all synonymous variants scores was 1 and nonsense variants 0.

For the abundance-based MAVE in HEK293T cells, barcodes were mapped from eight FASTQ files, representing the four GFP:mCherry ratio sort bins for each of the two replicate VAMP-seq experiments, with 1 being the lowest value ratio bin and 4 being the highest. Barcodes were filtered so that at a count of at least ten was observed within each sample. The counts for each barcode were collapsed to counts per variant, and then the frequency of each variant was calculated by dividing its count by the sum of all counts in that sample. Using the frequencies, a weighted average abundance score was calculated as reported previously^20^. Briefly, for each variant frequencies in bin 1 were multiplied by 0.25, bin 2 by 0.5, bin 3 by 0.75, and bin 4 by 1 and summed, then divided by the unweighted sum of that variant’s counts across the bins. The weighted averages were then normalized by scaling so the average of all synonymous variants scores was 1 and nonsense variants 0.

To compute a functional score for each variant, we selected the lower of the activity or abundance score, except for variants where there was no activity score; variants without activity scores only received a functional score if their abundance score was low (less than z-score of -3 from the average synonymous variant abundance score), indicating a likely loss of function due to low abundance. We re-normalized functional scores so the average functional score of all synonymous variants was 1 and nonsense variants 0.

### Structure mapping and analysis

Functional domains of G6PD were annotated based on previous reports^18,22^. Amino acid conservation between *S. cerevisiae* Zwf1 (CAA96146.1) and *H. sapiens* G6PD (QZA86926.1) was determined using the Sequence Manipulation Suite^42^, with default “similar” amino acid groupings of GAVLI, FYW, CM, ST, KRH, DENQ, P. Structure mapping was performed using Genomics 2 Proteins (G2P)^43^ on G6PD dimer structure 7UAG (https://doi.org/10.2210/pdb7UAG/pdb).

### Phenotype and clinical data

Phenotype data was curated previously and deposited at https://github.com/reneegeck/G6PDcat ^6^. Reports published through September 25, 2025 were considered in this literature analysis. For association of variants with nondeficiency, at least three individuals with over 60% normal activity were reported. Variants with any reported association with G6PD deficiency were categorized as “deficient;” they were subcategorized if AHA or favism were reported, or if at least 25% of cases reported had CNSHA. WHO classes were assigned as described^10^; briefly, variants with activity measurements in at least three unrelated individuals were class C if median activity was over 60%, class B if under 45%, and class A if under 20% and associated with CNSHA. ClinVar variant interpretations for contribution to G6PD deficiency were accessed September 25, 2025^7^.

### Variant effect predictors

REVEL scores for ENST00000393564 were downloaded from https://sites.google.com/site/revelgenomics/downloads and categorized by level of evidence described previously^44,45^. Alpha Missense scores and categories on ENST00000393564.6 were downloaded from https://alphamissense.hegelab.org/ ^46^.

### Level of evidence calibration

Score cutoffs were determined using the distribution of synonymous variant scores. A z-score of -1 was used as the minimum functional score for benign evidence, and -3 as the maximum for pathogenic evidence. ClinVar variant interpretations used for calibration were accessed September 25, 2025^7^. OddsPath was computed^27^ and translated to evidence strength thresholds using prior of 0.1. The OddsPath for benign and likely benign variants having functional scores over a z-score of -1 was 0.1579, translating to moderate evidence; the OddsPath for pathogenic and likely pathogenic variants having functional scores below a z-score of -3 was 51.01, translating to strong evidence.

### Data and resource availability

PacBio and Illumina sequencing data is available under SRA BioProject PRJNA1295945 and at https://data.igvf.org/analysis-sets/IGVFDS1322SJZB/. CountESS is available at https://zenodo.org/records/16357955. Code for analysis and generating figures is available at https://github.com/reneegeck/DunhamLab/tree/main/Geck2025_G6PD_MultiplexedFunctionalAs <u>say</u>. Strains and plasmids are available upon request.

## Acknowledgements

We thank Noah Fredstrom and Joe Armstrong for assistance with Illumina sequencing and Katy Munson for PacBio sequencing; Choli (Charlie) Lee for flow cytometry support; Gabrielle Ferra and Jochen Weile for assistance running Pacybara; Nick Popp for advice and assistance in analyses; Sandra Pennington, Scarlett Counihan, and Owen Burris for yeast media and reagent support; and Mary Relling, Jan Abkowitz, and members of the Dunham lab, Fowler lab, and ClinGen G6PD Variant Curation Expert Panel for feedback.

---

**Figure S1.**
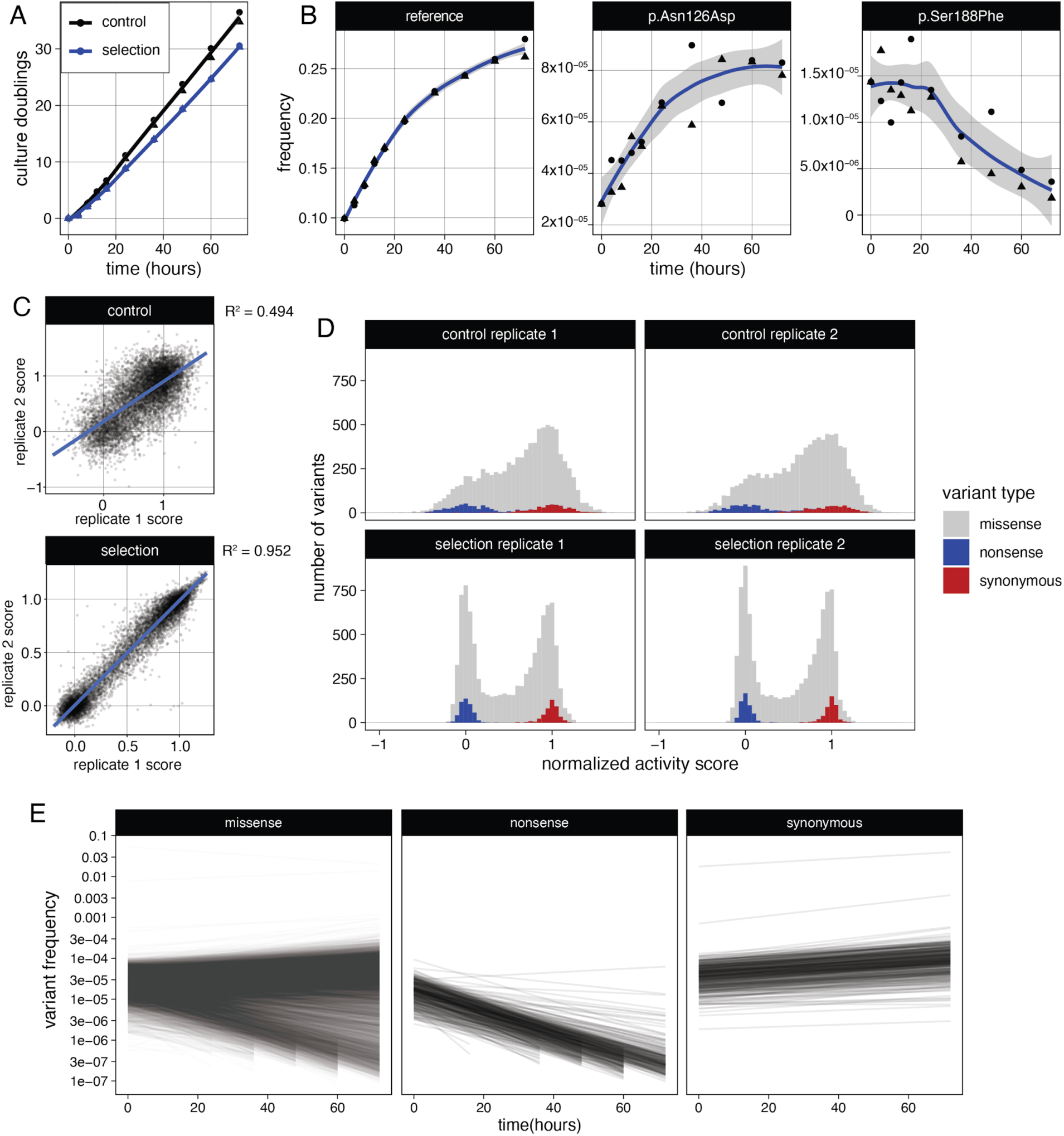
(A) Population doublings in the turbidostats, calculated from pump time. Shapes denote replicates. (B) Frequency of selected reference and well-characterised G6PD variants (G6PD A: p.Asn126Asp, and G6PD Mediterranean: p.Ser188Phe) within the selection turbidostats over time. Shapes denote replicates. (C) Normalized activity score correlation between replicates. (D) Distribution of variants by normalized activity scores. (E) Frequency of each variant over time in selection. Line represents linear fit to both selection replicates.

**Figure S2.**
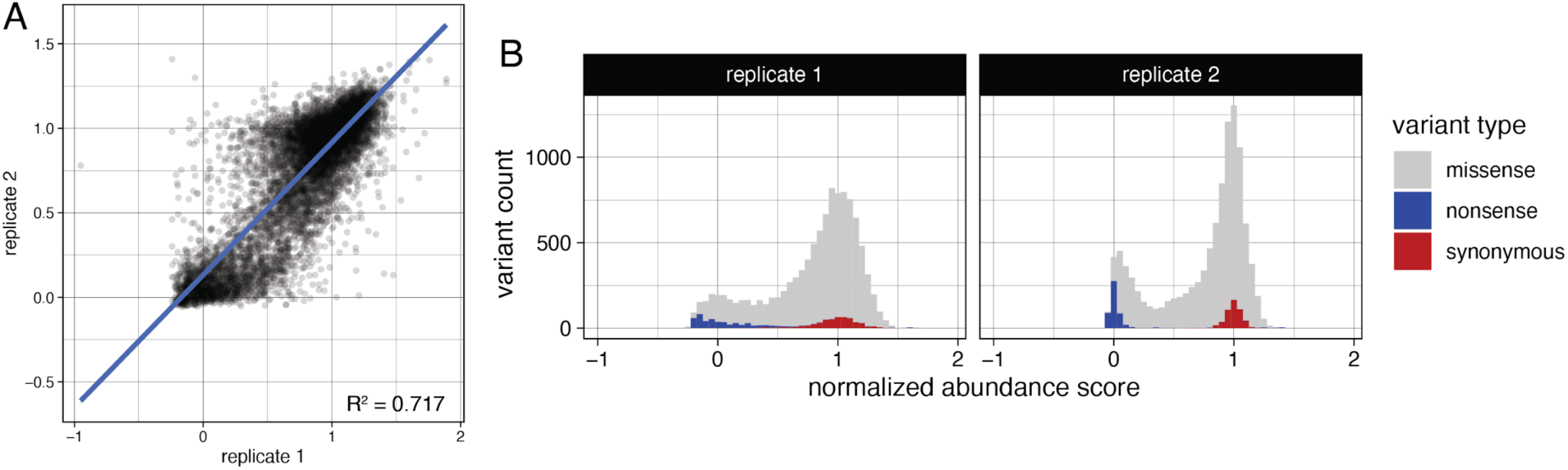
(A) Normalized abundance score correlation between replicates. (B) Distribution of variants by normalized abundance scores.

**Figure S3.**
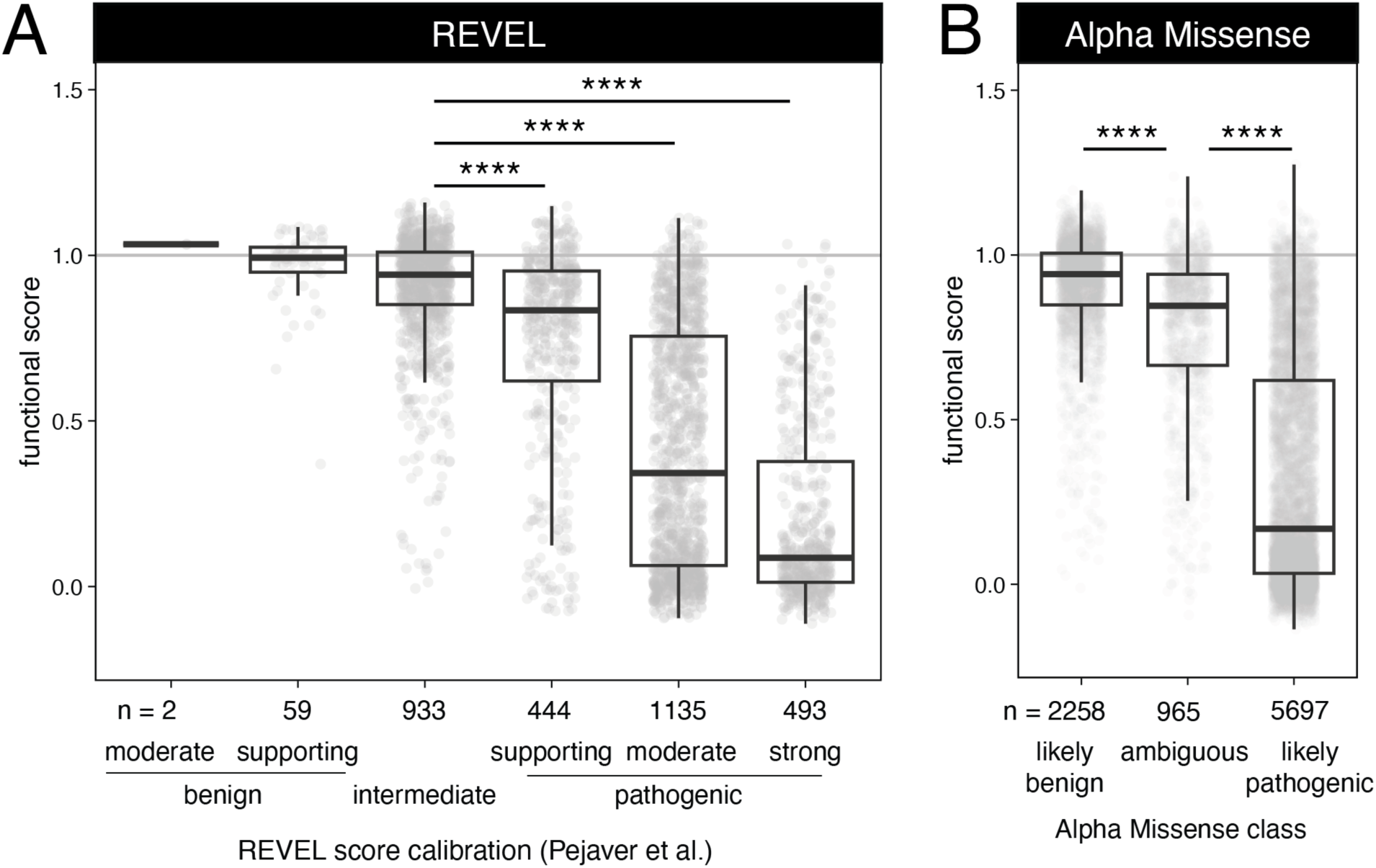
(A) *G6PD* variant functional scores compared to scores from VEPs REVEL and (B) Alpha Missense. Significance from by ANOVA with Tukey’s HSD, only showing statistics for difference compared to intermediate or ambiguous class; ****, p<0.0001.

